# Modified TCA/acetone precipitation of plant proteins for proteomic analysis

**DOI:** 10.1101/382317

**Authors:** Liangjie Niu, Hang Zhang, Zhaokun Wu, Yibo Wang, Hui Liu, Xiaolin Wu, Wei Wang

## Abstract

Protein extracts obtained from cells or tissues often require removal of interfering substances for the preparation of high-quality protein samples in proteomic analysis. A number of protein extraction methods have been applied to various biological samples. TCA/acetone precipitation and phenol extraction, a common method of protein extraction, is thought to minimize protein degradation and activity of proteases as well as reduce contaminants like salts and polyphenols. However, the TCA/acetone precipitation method relies on the complete pulverization and repeated rinsing of tissue powder to remove the interfering substances, which is laborious and time-consuming. In addition, by prolonged incubation in TCA/acetone, the precipitated proteins are more difficult to re-dissolve. We have described a modified method of TCA/acetone precipitation of plant proteins for proteomic analysis. Proteins of cells or tissues were extracted using SDS-containing buffer, precipitated with equal volume of 20% TCA/acetone, and washed with acetone. Compared to classical TCA/acetone precipitation and simple acetone precipitation, this protocol generates comparable yields, spot numbers, and proteome profiling, but takes less time (ca. 45 min), thus avoiding excess protein modification and degradation after extended-period incubation in TCA/acetone or acetone. The modified TCA/acetone precipitation method is simple, fast, and suitable for proteomic analysis of various plant tissues in proteomic analysis.

## Background

Protein extracts obtained from cells or tissues often contain interfering substances, which must be removed for preparing high-quality protein samples [1]. In particular, plant tissues contain a diverse group of secondary compounds, such as phenolics, lipids, pigments, organic acids, and carbohydrates, which greatly interfere with protein extraction and proteomic analysis [2]. Sample quality is critical for the coverage, reliability, and throughput of proteomic analysis; protein extraction in proteomics remains a challenge, even though advanced detection approaches (especially LC-MS/MS) can greatly enhance the sensitivity and reliability of protein identification. In fact, protein extraction methods shape much of the extracted proteomes [3].

A protein extraction protocol that can be universally applied to various biological samples with minimal optimization is essential in current proteomics. A number of methods are available for concentrating dilute protein solutions and simultaneously removing interfering substances, e.g., TCA/acetone precipitation and phenol extraction [9-11]. TCA/acetone precipitation is a common method for precipitation and concentration of total proteins, which was initially developed by Damerval et al. [12] and later modified by other workers for use in various tissues [10, 13, 14].

TCA/acetone precipitation is thought to minimize protein degradation and activity of proteases as well as reduce contaminants such as salts or polyphenols [15]. During acetone/TCA precipitation, organic-soluble substances are rinsed out, leaving proteins and other insoluble substances in the precipitate, and proteins are extracted using a buffer of choice [2, 5, 9]. The success of the TCA/acetone precipitation method is based on the complete pulverization and repeated rinsing of tissue powder to remove the interfering substances, which is a laborious and time-consuming process. However, prolonged incubation of tissue powder in TCA/acetone may lead to the modification of proteins by acetone, and the proportion of modified peptide increases over time, thus affecting the outcome of MS/MS analysis [16]. Moreover, long exposure to the acidic pH in TCA/acetone probably causes protein degradation [6,17]. Alternatively, protein extracts can be precipitated using aqueous 10% TCA [18], but TCA precipitated proteins are more difficult to dissolve and require the use NaOH to increase their solubilization [19]. Therefore, aqueous TCA precipitation, like TCA/acetone precipitation, is not commonly used in proteomic analysis.

There are other alternatives to the use of TCA/acetone for protein precipitation in proteomics, e.g., acetone precipitation, acetonitrile/trifluoroacetic acid precipitation, and methanol/chloroform precipitation. However, these methods have some limitations. Acetone precipitation needs at least a 4:1 ratio of acetone to the aqueous protein solution [20], which is not convenient for precipitating a large volume of protein extract, especially in Eppendorf tubes. Methanol/chloroform precipitation was developed for protein recovery from a small volume (e.g., 0.1 ml) of dilute solution [21], and acetonitrile precipitation is commonly used for recovery of peptides from trypsin-digested gel pieces for mass spectrometry [22].

To overcome the limitations of simple acetone precipitation, aqueous TCA precipitation, and TCA/acetone precipitation, we report, herein, a modified, rapid method of TCA/acetone precipitation of plant proteins for proteomic analysis. We systematically compared the modified TCA/acetone precipitation, classical TCA/acetone precipitation, and simple acetone precipitation methods with respect to protein yields and proteome profiles and analyzed the coefficient of variation of each spot in 2DE maps from three independent experiments.

## Methods

### Materials

Maize (*Zea mays* L. cv Zhengdan 958) was used as the experimental material. To sample maize embryos, mature seeds were soaked in water for 2 h to soften seed coats and endosperms, and the embryos were manually dissected and used for protein extraction. For maize mesocotyl sampling, mature seeds were sterilized with 0.1% sodium hypochlorite and were germinated on moistened filter paper for 3d (28°C) to excise the mesocotyl (ca. 2.0-2.5 cm) for protein extraction. For maize leaf and root sampling, dark-germinated seedlings were then cultured in Hogland’s nutrient solution in a light chamber (day 28°C/night 22°C, relative humidity 75%) under 400 μ mol m^-2^ s^-1^ photosynthetically active radiation with a 14/10 h (day/night) cycle for two weeks [23]. The fully-expanded 3rd leaves and 1 cm-long root tips were collected for protein extraction.

### Modified TCA/Acetone precipitation of proteins

The modified protocol is designed for running in Eppendorf tubes within 1 h and can be reliably adapted to big volumes. It includes protein extraction, precipitation, and dissolving. The detailed steps are given in Fig. 1. Maize tissues were used for evaluating the protocol. The organic solvents used here, including acetone, 80% acetone, and 20% TCA/acetone, were pre-cooled at −20°C and were supplemented with 5 mM dithiothreitol (DTT) before use. The subsequent steps were carried out at 4°C unless otherwise indicated.

**Fig 1.**
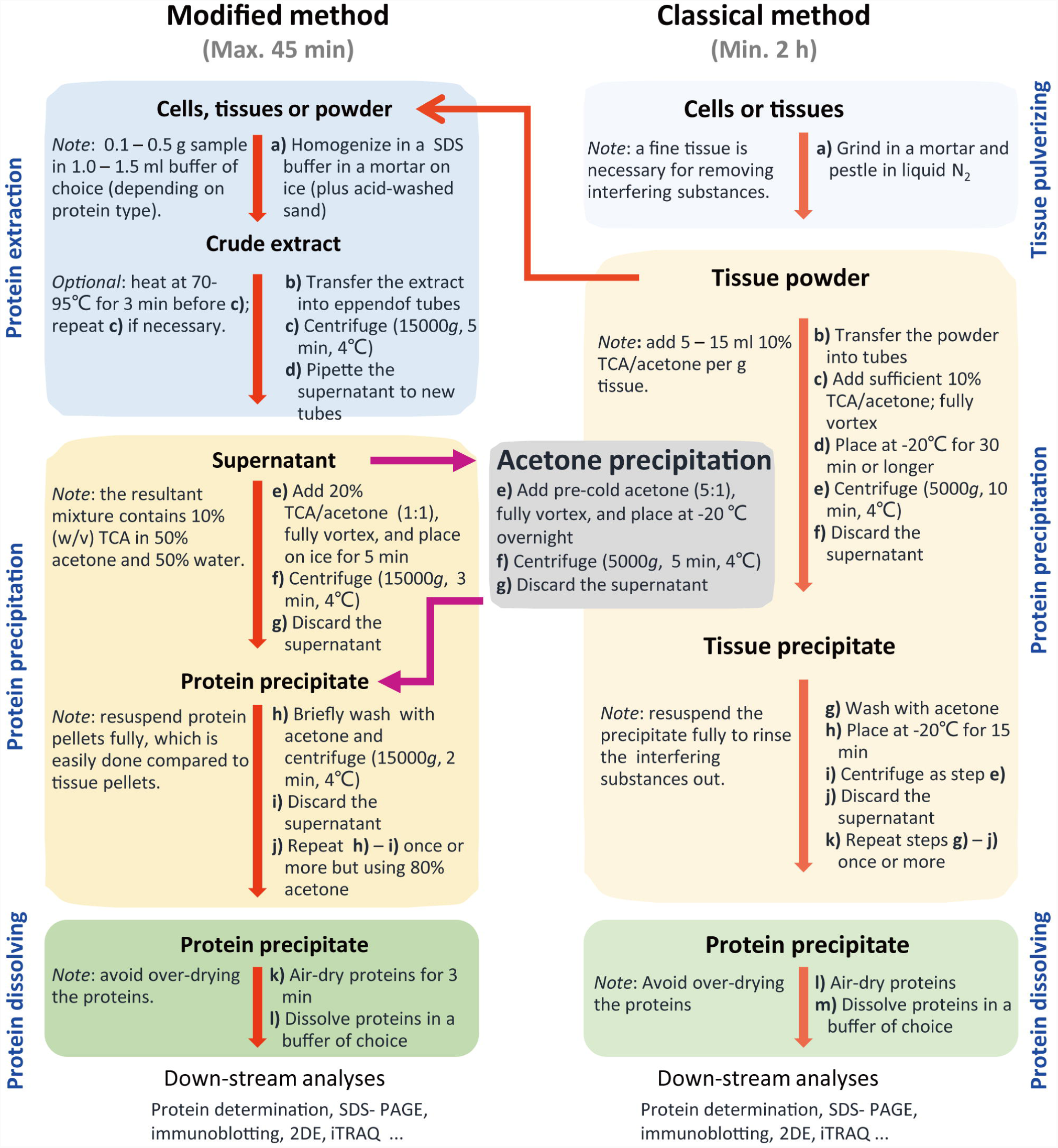
Comparison between the steps of the modified TCA/acetone precipitation, the classical TCA/acetone precipitation, and acetone precipitation methods. The SDS extraction buffer contained 1% (w/v) SDS, 0.1 M Tris-HCl (pH 6.8), 2 mM EDTA-Na_2_, 20 mM DTT, and 2 mM PMSF (added before use). All organic solvents were pre-chilled at −20°C and contained 5 mM DTT (added before use).

#### Protein extraction

Maize embryos (0.2g), leaves (0.4g), mesocotyl (0.4 g) and roots (0.4g) were homogenized in a pre-cooled mortar (interior diameter 5 cm) on ice in 2.0 ml of the extraction solution containing 1% SDS, 0.1 M Tris-HCl (pH 6.8), 2 mM EDTA-Na_2_, 20 mM DTT, and 2 mM PMSF (added before use). The homogenate was transferred into Eppendorf tubes and centrifuged at 15,000 g for 5 min. Then, the supernatant (protein extract) was pipetted into fresh tubes.

#### Protein precipitation

20% cold TCA/acetone was added to the protein extract (1:1, v/v, with a final 10% TCA/50% acetone), the mixture was placed on ice for 5 min, centrifuged at 15,000 g for 3 min and the supernatant was discarded. The protein precipitate was washed with 80% acetone, followed by centrifugation as above. The wash step was repeated once or more.

#### Protein dissolving

The protein precipitates were air-dried for a short duration (1-3 min) and dissolved in a buffer of choice for protein analysis. Notably, the precipitates should not be over-dried as this makes it more difficult to resolubilize them.

### TCA/ acetone precipitation

TCA/acetone precipitation was done exactly as previously described [10]. Briefly, plant tissues were pulverized to a fine powder in a mortar in liquid N_2_. The powder was suspended in 10% TCA/acetone and kept at –20°C overnight. Then, the samples were centrifuged for 30 min at 5,000 g at 4°C. The resultant pellets were rinsed with cold acetone twice, and each step involved a centrifugation for 10 min at 5,000 g at 4°C. The protein precipitates were air-dried for a short duration (1-3 min) and dissolved in a buffer of choice for protein analysis.

### Acetone precipitation

One-step acetone precipitation was performed as described recently [20]. Protein extracts were precipitated with 6 volumes of cold acetone and kept at −20°C overnight, followed by two pellet-washing steps, each with cold acetone. Protein pellets were collected by centrifugation at 10,000 g at 4°C for 30 min, air-dried for 15 min in the ice box, and dissolved in a buffer of choice for protein analysis.

### Protein assay

For SDS-PAGE, protein precipitates were dissolved in a SDS-containing buffer (0.5% SDS, 50mM Tris-HCl, pH 6.8, and 20 mM DTT). Protein concentration was determined using the Bio-Rad Bradford assay kit (Bio-Rad, Hercules, CA) [24], but performed on a micro scale, i.e., 10 μl of standard or sample solution was mixed with 1.0 ml of diluted dye solution. In this way, the final concentration of SDS in the mixture was 0.005%, which was compatible with the Bradford assay. Prior to SDS-PAGE, protein extracts were mixed with appropriate volume 4 x SDS sample buffer [25]. For 2DE, protein precipitates were dissolved in the 2DE rehydration solution without IPG buffer to avoid its interference as we described before [26], and protein concentrations were determined by the Bradford Assay. Subsequently, the IPG buffer was supplemented into protein samples to a concentration of 0.5%.

### SDS-PAGE

SDS-PAGE was run using 12.5% gel [27] and protein was visualized using Coomassie brilliant blue (CBB) G250.

### 2-DE and MS/MS

Isoelectric focusing (IEF) was performed using 11-cm linear IPG strips (pH 4–7, Bio-Rad). Approximately 600 μg of proteins in 200 μl of the rehydration solution was loaded by passive rehydration with the PROTEAN IEF system (Bio-Rad) for 12 h at 20°C. IEF and subsequent SDS-PAGE, and gel staining were performed as previously described [27]. Digital 2-DE images were processed and analyzed using PDQUEST 8.0 software (Bio-Rad). Protein samples were analyzed by 2DE in three biological replicates.

The spots with at least 2-fold quantitative variations in abundance among maize embryos, leaves, and roots by two methods, respectively, were selected for mass spectrometry (MS) analysis. One-way ANOVA was performed based on three biological replications.

The selected protein spots were extracted, digested, and analyzed by the MALDI-TOF/TOF analyzer (AB SCIEX TOF/TOF-5800, USA) as described previously [28]. MALDI-TOF/TOF spectra were acquired in the positive ion mode and automatically submitted to Mascot 2.2 (http://www.matrixscience.com) for identification against NCBInr database (version Sept 29, 2018; species, *Zea mays*, 719230 sequences). Only significant scores defined by Mascot probability analysis greater than “identity” were considered for assigning protein identity. All the positive protein identification scores were significant (p<0.05).

### Bioinformatics Analysis

The unidentified proteins were searched by BLAST using Universal Protein (http://www.uniprot.org/) or National Center for Biotechnology Information (http://www.ncbi.nlm.nih.gov/). Grand average of hydropathicity (GRAVY) analyses were performed using ProtParam tool (http://web.expasy.org/protparam/). Subcellular localization information was predicted using the online predictor Plant-mPLoc (http://www.csbio.sjtu.edu.cn/bioinf/plant-multi/).

## Results

### The development of the modified TCA and acetone precipitation

Rather than preparing tissue powder by extensive TCA/acetone rinsing using the classical method, we directly extracted proteins in cells or tissues and then precipitated proteins in the extracts with equal volume of 20% TCA/acetone. Various aqueous buffers can be used for protein extraction, but the composition of the extraction buffers will greatly affect the profiles of the extracted proteome. The Laemmli’s SDS buffer [25] was used for protein extraction, because SDS-containing buffers can enhance protein extraction and solubility, especially under heating.

The ratio of TCA and acetone was optimized in initial tests (Fig. 2). We compared the effect of 10% (w/v) TCA in various concentrations (0-80%, v/v) of aqueous acetone and the effect of various TCA concentrations in 50% aqueous acetone on protein extraction and separation. The presence of TCA could improve protein resolution by SDS-PAGE, but no significant differences were observed in TCA concentrations of 5-20% (Fig. 2A). At a fixed 10% TCA, protein patterns were quite similar with different acetone ratios, but the background was clearer with increased acetone ratio (Fig. 2B). Based on the quality of protein gels, we chose the low ratio combination of 10% TCA/50% acetone (final concentration) for further testing.

**Fig 2.**
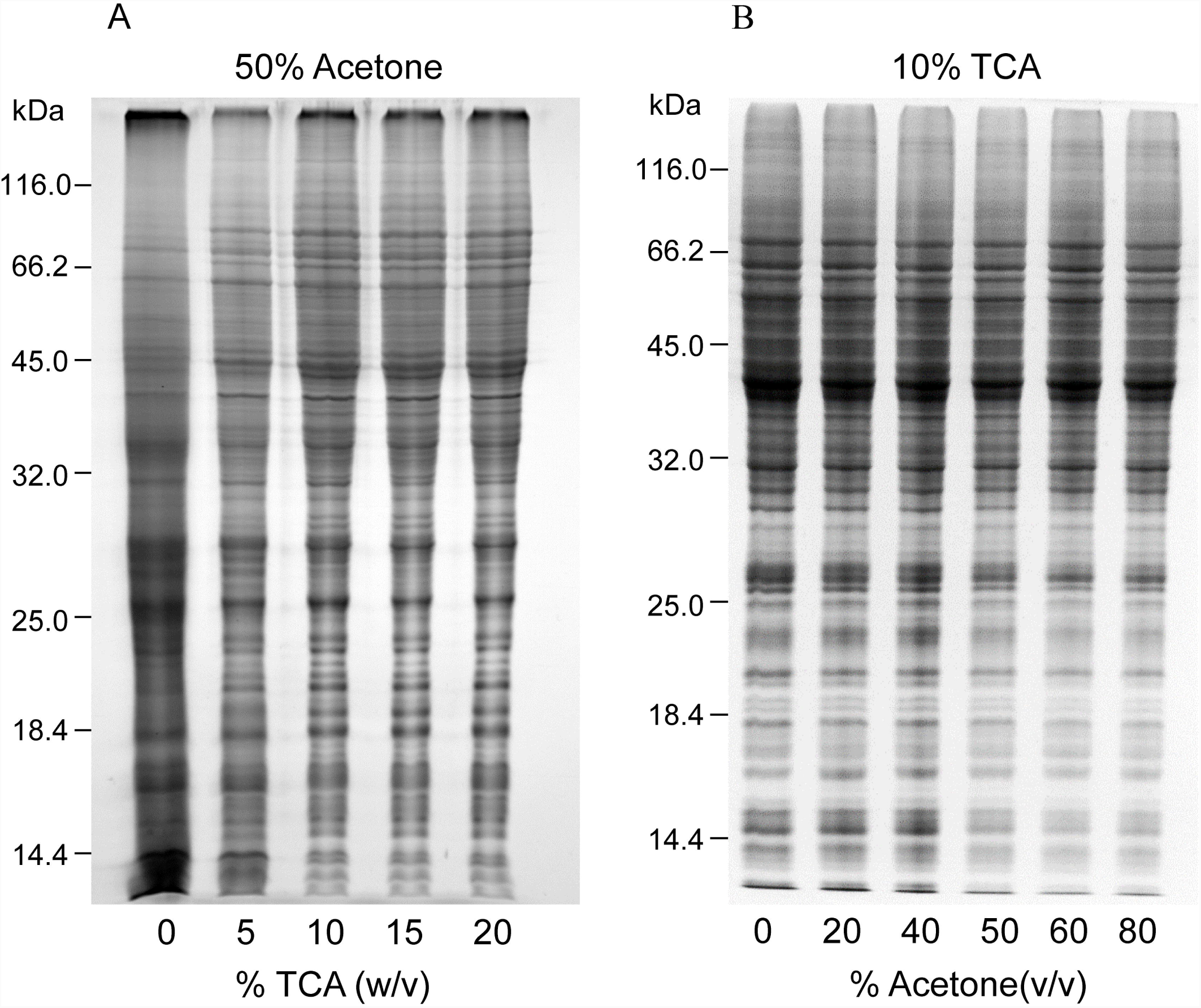
Optimization of TCA/acetone ratio used in the modified method. Equal amounts (ca. 30 μg) of maize root proteins were analyzed by SDS-PAGE (12.5% resolving gel). Protein was stained using CBB. A, final acetone concentration was 50% (v/v), but TCA concentration varied from 0-20% (w/v) in the aqueous mixture. B, final TCA concentration was 10% (w/v), but acetone concentration varied from 0-80% (v/v) in the aqueous mixture.

As opposed to aqueous TCA precipitation, proteins precipitated by 10% TCA/50% acetone are easy to dissolve. After incubation on ice for 10 min, protein precipitates were recovered by centrifugation and washed with 80% acetone thrice to remove residual TCA in the precipitated protein. Finally, the air-dried protein precipitates were dissolved in a buffer of choice for SDS-PAGE, IEF, or iTRAQ analysis.

### Evaluation of the modified method

First, we made a comprehensive comparison of protein yields and resolution in 2-DE by the modified and classical methods. The protein yields were slightly higher in the modified method but did not differ significantly from the classical method (p<0.05). Similar results were obtained by comparison of spot numbers in 2D gels (Table 1). Compared to other separation methods (e.g., SDS-PAGE, LC), 2DE analysis can visibly display the quality of protein samples.

**Table 1.** Comparison of protein yield and spot number in 2DE between the two methods.

| Maize tissues | The modified method |  | The classical method |  |
| --- | --- | --- | --- | --- |
|  | Yield <sup>a</sup> | Spot No <sup>b</sup> | Yield <sup>a</sup> | Spot No <sup>b</sup> |
| Roots | 4.82±0.07 | 738±11 | 4.70±0.08 | 743±10 |
| Leaves | 4.13±0.11 | 310±9 | 4.06±0.23 | 314±4 |
| Embryos | 5.80±0.13 | 314±12 | 5.56±0.38 | 304±18 |

Notes: a, Protein yields are expressed as µg/mg fresh weight. b, Average spot number detected in 2D gels using PDQUEST software (version 8.0, Bio-Rad). All data were from at least three independent experiments. The corresponding data from the two methods did not differ significantly (p<0.05) according to t-test.

For all materials tested, 2D gels obtained with the two methods were generally comparable regarding the number, abundance, and distribution of protein spots, without profound deviations (Fig3, Fig S1-S3); however, several protein spots exhibited at least 2-fold differences in abundance. For example, spots 1, 2, and 4 were more abundant in maize roots by the modified method, whereas the classical method resulted in more abundant spot 3. Of the 11 differential abundance proteins (DAPs) selected for MS/MS identification, nine were identified with MS/MS analysis (Table 2). In particular, the modified method selectively depleted globulin-1 in maize embryos (Fig. 3). Our recent studies showed that globulin-1 (also known as vicilin) is the most abundant storage protein in maize embryos [11, 29], and selective depletion of globulin-1 improved proteome profiling of maize embryos [11].

**Table 2.**
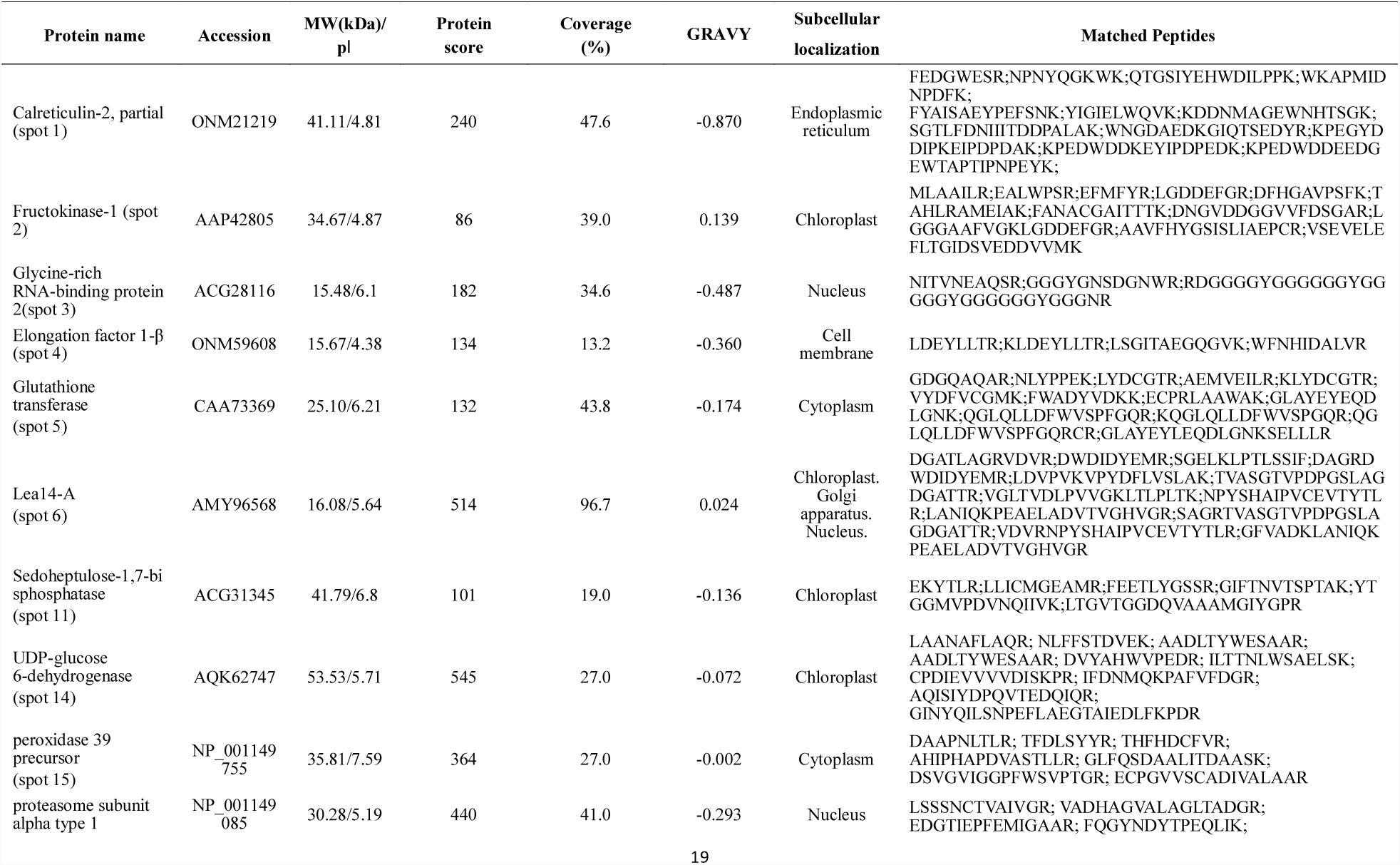
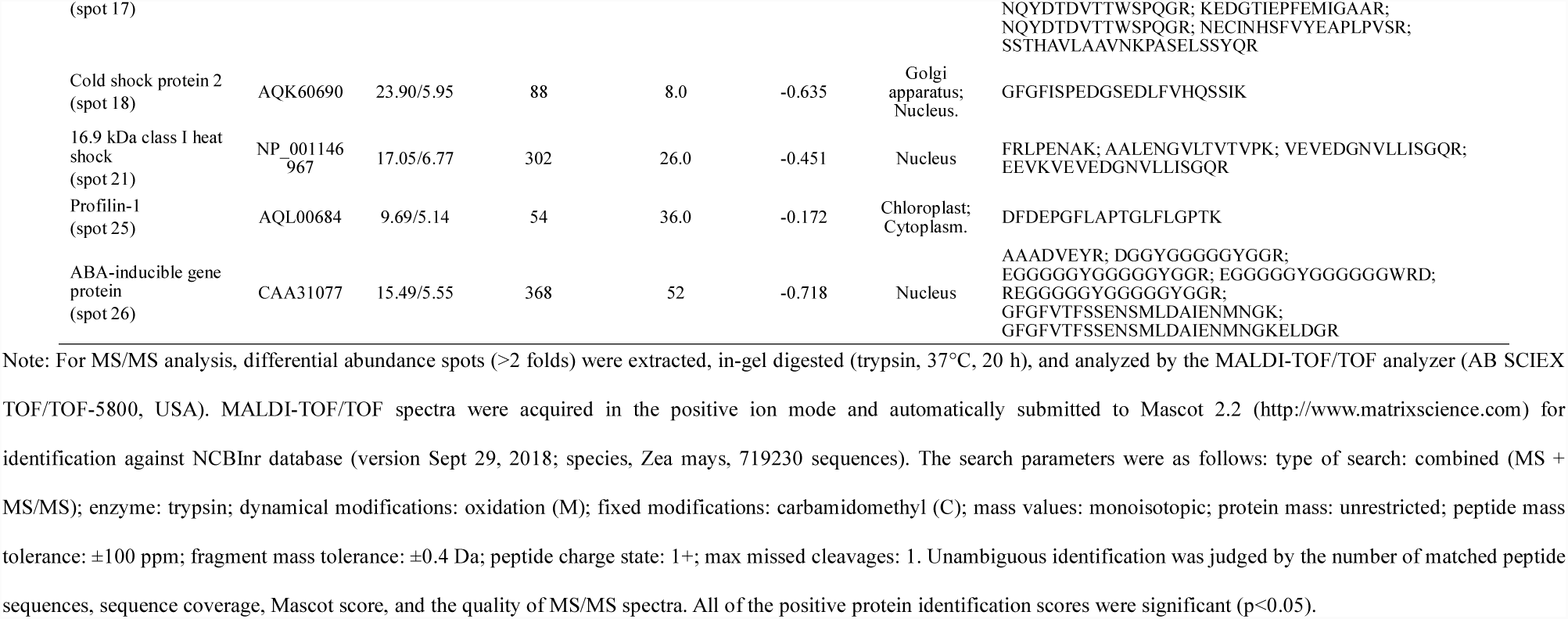
The identification of the differential extracted proteins in maize using the two methods

**Fig 3.**
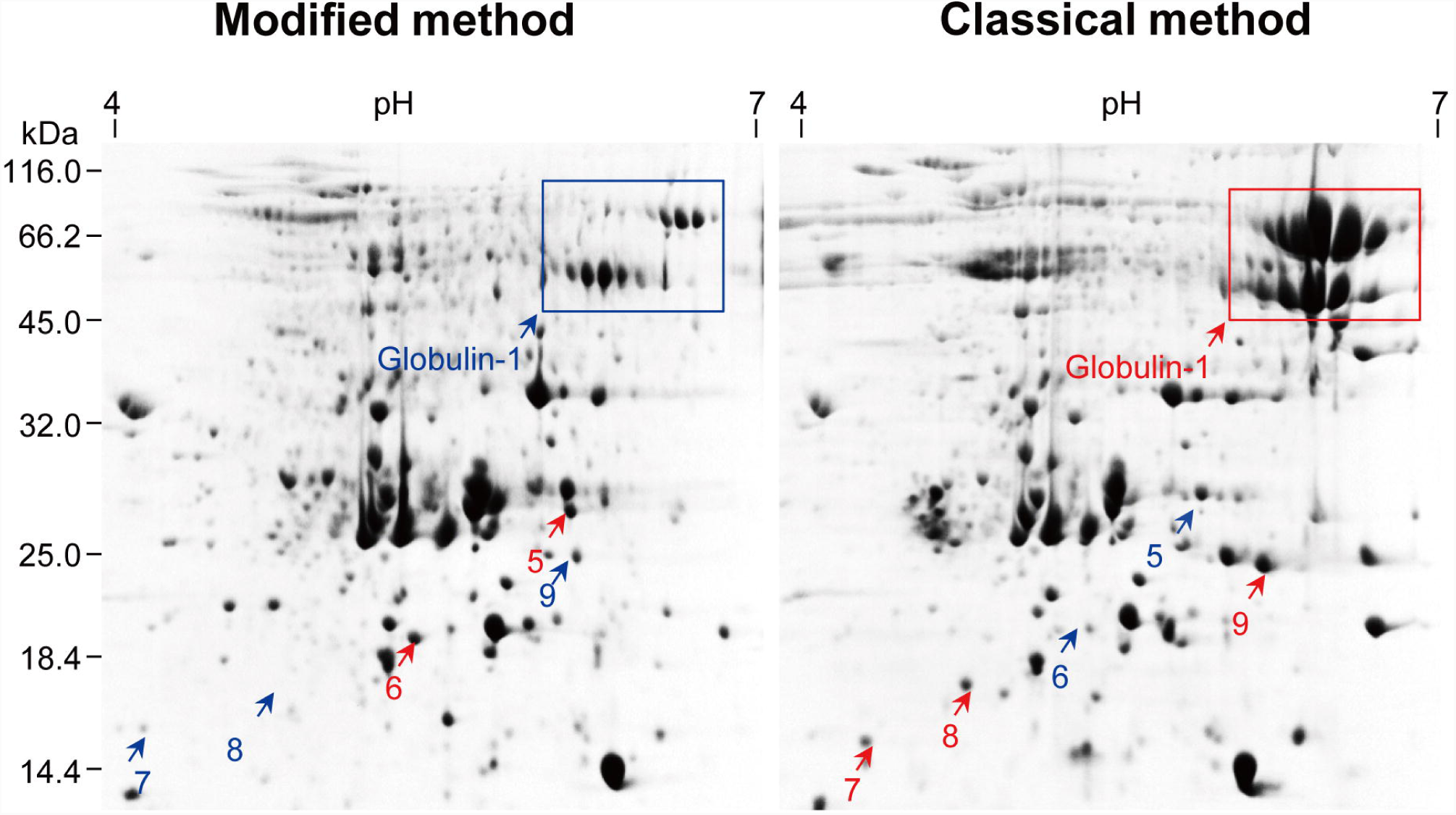
Comparison of 2DE protein profiles of maize embryo proteins extracted using two methods. *Left panel*: the modified TCA/acetone precipitation. *Right panel*: the classical TCA/acetone precipitation. Spots with increased abundance are indicated in red. About 800 µg of proteins were resolved in pH 4-7 (linear) strip by IEF and then in 12.5% gel by SDS-PAGE. Proteins were visualized using CBB.

Second, we analyzed each of the spot variations in 2DE gels obtained with the two methods from three independent replicates Fig 4, Fig S4, Table S1). Particularly, for 2DE gels of maize mesocotyls, 553, 558, and 605 colloidal CBB stained spots were detected in three 2DE maps, respectively, with 437 spots in common. Spot-to-spot comparison revealed a reproducibility of 72-79% (matched spots/total spots ratio). Comparable results were obtained for 2DE gels of maize embryos (Fig. 3, Table S2). Undeniably, there was a substantial variation in abundance among spots in three independent replicates, which is an inherent drawback of common 2DE.

**Fig 4.**
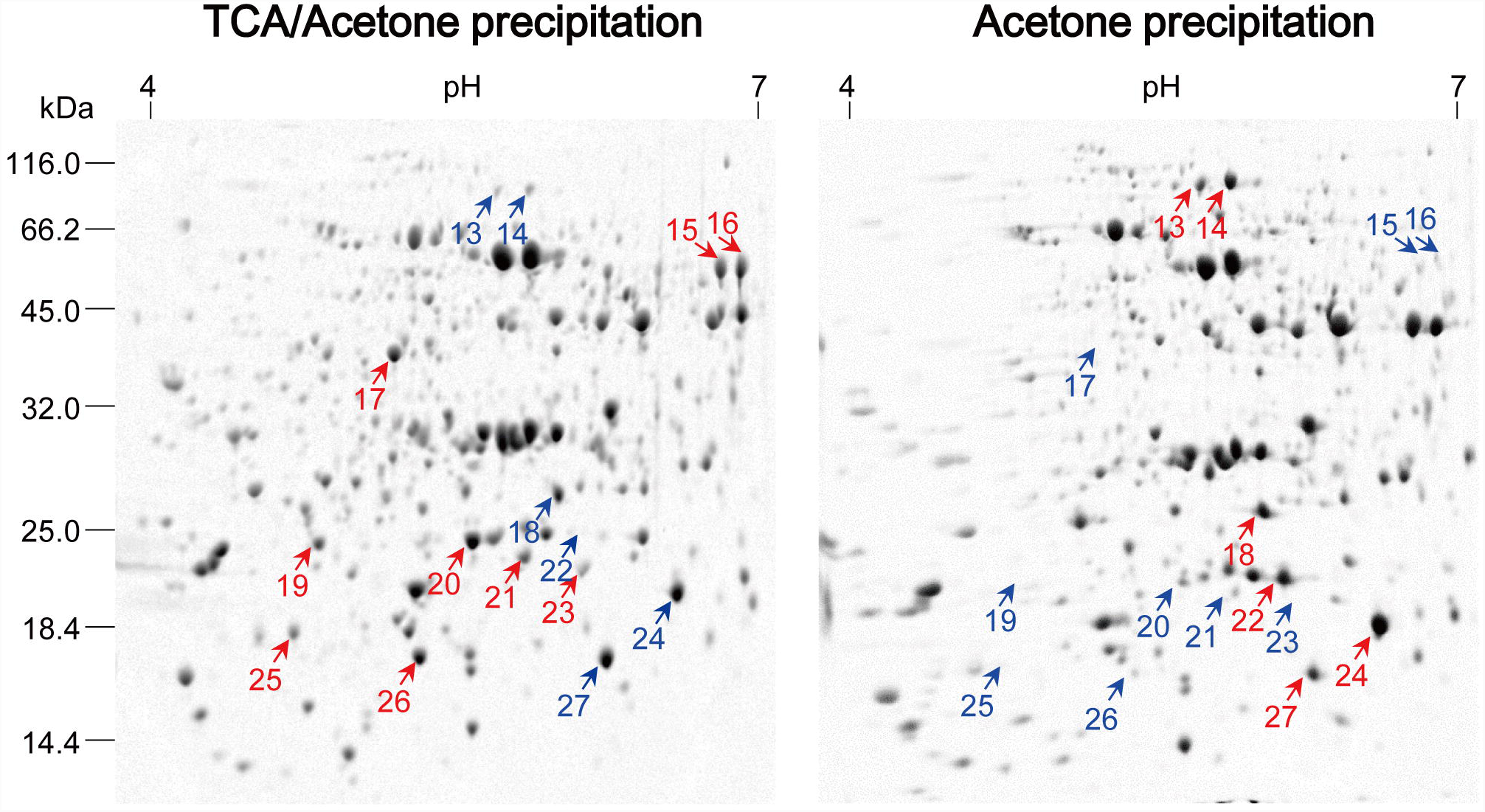
Comparison of 2DE profiles of maize mesocotyl proteins extracted using two methods. Another two independent experiments were shown in Fig S4. Left panel: the modified TCA/acetone precipitation. Right panel: acetone precipitation. About 800 µg of proteins were resolved in pH 4-7 (linear) strip by IEF and then in 12.5% gel by SDS-PAGE. Protein was visualized using colloidal CBB.

Finally, we compared the modified acetone/TCA precipitation and simple acetone precipitation methods (Fig 4, Fig S4). Overall, the former produced good 2DE maps. Obviously, some spots were preferably extractable to the extraction method, but more spots were lost after simple acetone precipitation, especially high-mass spots in acidic regions. Some of proteins of interest were subjected to MS/MS identification (Table 2). Though simple acetone precipitation worked well for some cell materials [20]. Many previous studies indicated that simple acetone precipitation precludes production of good 2DE maps due to the presence of high levels of interfering substances in plant materials. A recent research reported that protein loss is believed to be an inevitable consequence of acetone precipitation of proteome extracts [30].

In addition, it is worthwhile to note that aqueous TCA precipitation can cause severely denatured proteins that are very difficult to dissolve; hence, this method is rarely used in proteomic analysis. Thus, we did not compare aqueous TCA precipitation with the modified method in the present study.

## Discussion

The classical TCA/acetone precipitation method applies a strategy of removal of interfering substances before protein extraction, involving incubation for extended periods (from 45 min to overnight) in TCA/acetone and between the rinsing steps [10,12,15]. These steps can lead to the modification of proteins by acetone [16] or possible protein degradation after long exposure to harsh TCA/acetone [6,18,30], thus affecting the outcome of MS/MS analysis.

In contrast, the modified method described here uses a strategy of removing interfering substances after protein extraction, taking less time and thereby avoiding protein modification by TCA/acetone, but producing similar or better results regarding protein yields, 2DE spot numbers, and proteome profiling. The resultant protein precipitates are easy to wash using acetone compared to tissue powder in the classical TCA/acetone precipitation method.

Previously, Wang et al. [28] observed that oil seed protein extraction uses 10% TCA/acetone, rather than aqueous TCA, because the former results in protein precipitates which are easily dissolved in SDS buffer or 2D rehydration buffer. The combination of TCA and acetone is more effective than either TCA or acetone alone to precipitate proteins [13,31]. It is noted that some proteins were preferentially extracted by the modified or the classical method, but the rationale remains open to question. Recently, we reported a chloroform-assisted phenol extraction method for depletion of abundant storage protein (globulin-1) in monocot seeds (maize) and in dicot (soybean and pea) seeds [8]. The modified method was highly efficient in depleting globulin-1, suggesting another application of the modified method in proteomic analysis.

In the modified method, final protein pellets were dissolved in the same 2DE buffer as in the classical method, so protein profiles are highly dependent on the extraction efficiency in the SDS buffer and precipitation efficiency by 20% TCA/acetone. We observed some significant, repeated differences in abundance of several DAPs between the two methods. There were specific DAPs associated with each method in different samples. However, the reason behind this phenomenon remains unclear. We tried to analyze the hydropathicity, physicochemical property, and subcellular compartments of these DAPs (Table 2); however, no definite conclusion could be drawn. Understandably, different extraction methods can produce protein profiles with substantial or subtle differences [32], but these inherent differences are difficult to explain, as discussed in a previous study [33]. It is important to note that protein loss is an inevitable consequence of solvent precipitation, even in the modified method, as observed in acetone precipitation of proteome extracts, including bacterial and mammalian cells [30].

To summarize, the greatest advantages of the modified method are its simplicity and fast. Despite its steps being similar to aqueous TCA precipitation, the modified method circumvents the drawback of aqueous TCA precipitation, namely, TCA-precipitated proteins being difficult to dissolve. Moreover, the modified method uses equal volume 20% TCA/acetone to precipitate proteins and can handle bigger volumes of protein extracts than acetone precipitation in a microtube. As the modified method precipitates proteins in aqueous extracts, it is expected to be universally applicable for various plant tissues in proteomic analysis.

## Supporting information

## Acknowledgements

Financial support for this study was provided by National Natural Science Foundation of China (http://www.nsfc.gov.cn/, grant no. 31771700) and the Program for Innovative Research Team (in Science and Technology) in University of Henan Province, china (http://www.haedu.gov.cn/, grant no. 15IRTSTHN015).

## Supporting Information Legends

**Fig S1.** Comparison of 2DE protein profiles of maize embryo proteins extracted using two methods. Shown were two independent experiments. *Left panel*: the modified TCA/acetone precipitation. *Right panel*: the classical TCA/acetone precipitation. Spots with increased abundance are indicated in red. About 800 µg of proteins were resolved in pH 4-7 (linear) strip by IEF and then in 12.5% gel by SDS-PAGE. Proteins were visualized using CBB.

**Fig S2.** Comparison of 2DE protein profiles of maize root proteins extracted using two methods. Shown were three independent experiments. Left panel: the modified TCA/acetone precipitation. Right panel: the classical TCA/acetone precipitation. Spots with increased abundance are indicated in red. About 800 µg of proteins were resolved in pH 4-7 (linear) strip by IEF and then in 12.5% gel by SDS-PAGE. Proteins were visualized using CBB.

**Fig S3.** Comparison of 2DE protein profiles of maize leaf proteins extracted using two methods. Shown were three independent experiments. Left panel: the modified TCA/acetone precipitation. Right panel: the classical TCA/acetone precipitation. Spots with increased abundance are indicated in red. About 800 µg of proteins were resolved in pH 4-7 (linear) strip by IEF and then in 12.5% gel by SDS-PAGE. Proteins were visualized using CBB.

**Fig S4.** Comparison of 2DE profiles of maize mesocotyl proteins extracted using two methods. Shown are two independent experiments. Left panel: the modified TCA/acetone precipitation. Right panel: acetone precipitation. About 800 µg of proteins were resolved in pH 4-7 (linear) strip by IEF and then in 12.5% gel by SDS-PAGE. Protein was visualized using colloidal CBB.

**Table S1.** Comparison of spot variations in 2DE gels of maize mesocotyl proteins extracted using the modified TCA/acetone precipitation methods in three independent replicates. 2DE gels were processed and analyzed using PDQuest.

**Table S2.** Comparison of spot variations in 2DE gels of maize embryo proteins extracted using the modified TCA/acetone precipitation methods in three independent replicates. 2DE gels were processed and analyzed using PDQuest.

