## Supplementary figures and images for "Modified TCA/acetone precipitation of plant proteins for proteomic analysis"

### Supplementary file 1

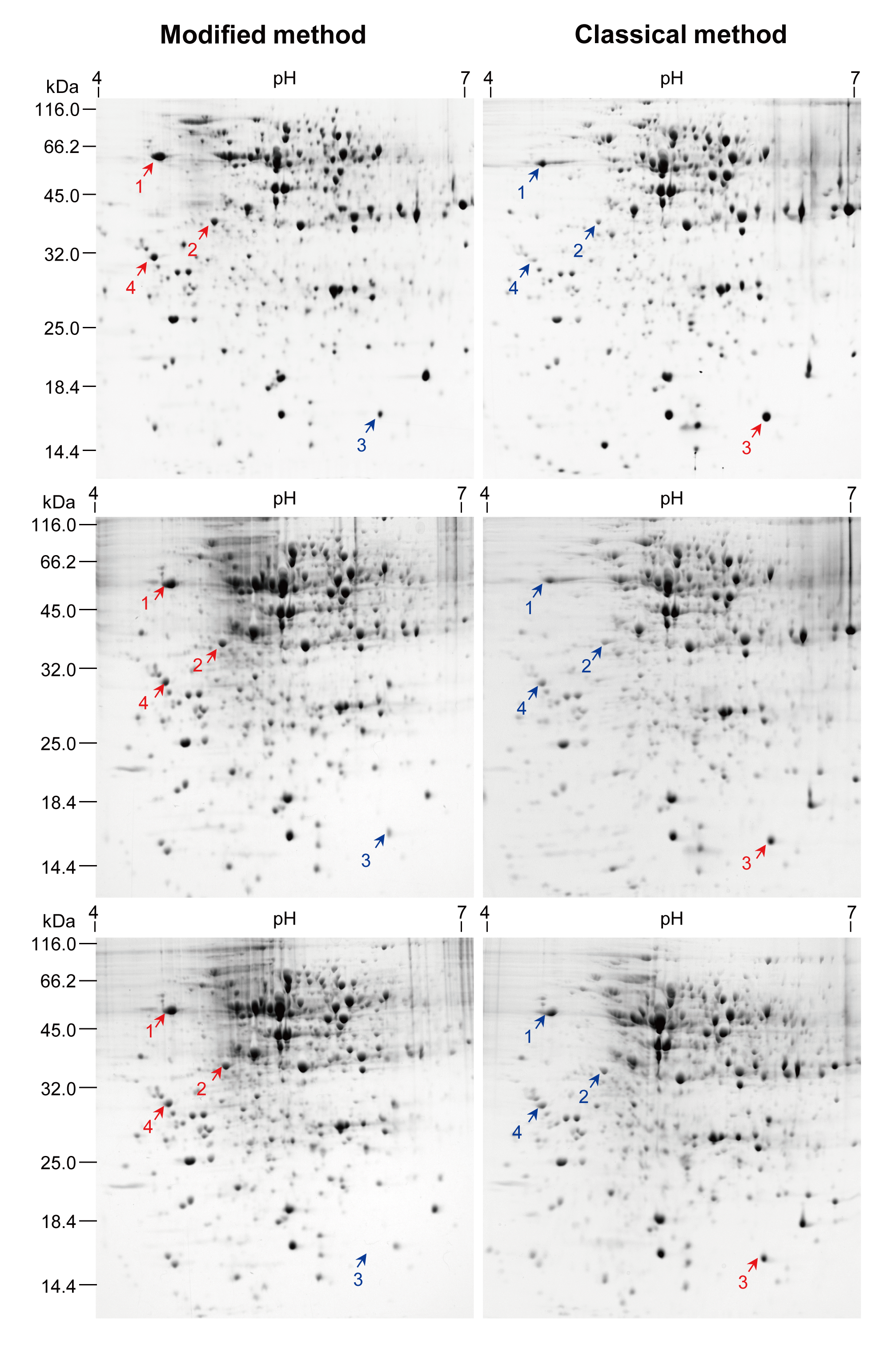
